# Phasic Dopamine Release Magnitude Tracks Individual Differences in Sensitization of Locomotor Response following a History of Nicotine Exposure

**DOI:** 10.1101/646844

**Authors:** Ashley M. Fennell, Elizabeth G. Pitts, Lacey L. Sexton, Mark J. Ferris

## Abstract

Smoking remains the primary cause of preventable death in the United States and smoking related illness costs more than $300 billion annually. Nicotine (the primary reinforcer in cigarettes) causes changes in behavior and neurochemistry that lead to increased probability of relapse. Given the role of mesolimbic dopamine projections in motivation, substance use disorder, and drug relapse, we examined the effect of repeated nicotine on rapid dopamine signals in the nucleus accumbens (NAc) of rats. Adult, male Sprague-Dawley rats were exposed to nicotine (0.2 or 0.4 mg/kg, subcutaneous) once daily for 7 days. On day 8, dopamine release and uptake dynamics, and their modulation by nicotinic receptor agonists and antagonists, were assessed using fast scan cyclic voltammetry in the NAc core. Nicotine exposure decreased electrically-stimulated dopamine release across a range of stimulation frequencies and decreased α6β2-containing nicotinic receptor control over dopamine release. Additionally, nicotine locomotor sensitization correlated with accumbal dopamine modulation by nicotine and mecamylamine. Taken together, our study suggests that repeated exposure to nicotine blunts dopamine release in the NAc core through changes in α6β2 modulation of dopamine release and individual differences in the sensitivity to this outcome may predict variation in behavioral models of vulnerability to substance use disorder.

## Introduction

Smoking tobacco is the number one cause of preventable death in the United States, with 480,000 individuals dying each year from cigarette use and second-hand smoke exposure (U.S. Department of Health and Human Services, 2014). Nicotine, the main reinforcer in tobacco, is a primary reinforcer that has been shown to support self-administration, increase and sensitize locomotor activity, and drive drug-seeking behavior (Stolerman & Shoaib, 1991; De Biasi & Dani, 2011). Additionally, nicotine enhances the reinforcing effects and incentive motivation of stimuli that accompany tobacco use (Caggiula et al., 2009).

Nicotinic acetylcholine receptors (nAChRs) are necessary for both the primary reinforcing and reinforcement enhancing effects of nicotine. Activation of nAChRs in the nucleus accumbens or in the VTA can directly increase dopamine release in the striatum (Cachope et al., 2012; Threlfell et al., 2012) and systemic nAChR antagonism decreases nicotine self-administration (Corrigall & Coen, 1989; DeNoble & Mele, 2006; Yoshimura et al., 2007). In addition, NAc nAChRs modulate dopamine release in a frequency dependent manner (Zhang & Sulzer, 2004). Dopamine neurons switch between tonic (∼4-5 Hz) and phasic (2-5 spikes at 20-100 Hz) patterns of firing during the presentation of reinforcers or reward-related cues (Waelti et al., 2001; Tobler et al., 2005; Marinelli & McCutcheon, 2014). Nicotine is thought to enhance the contrast between baseline firing and reward-related firing by decreasing dopamine release to tonic firing rates while increasing dopamine release to phasic firing patterns in the NAc (Rice & Cragg, 2004). This is hypothesized to enhance the salience of reward-related cues and play a role in the reinforcement enhancing effects of nicotine. Further supporting this hypothesis, systemic antagonism of nAChRs decreases nicotine-induced enhancement of reinforcers, although the brain regions necessary for this effect have not yet been established (Liu et al., 2007; Palmatier et al., 2009).

Repeated exposure to nicotine upregulates nAChRs in the striatum (see Gentry & Lukas, 2002; Penton & Lester, 2009). Repeated nicotine also alters nAChR modulation of dopamine in the striatum. Two studies found that chronic oral nicotine self-administration in mice decreases electrically-stimulated dopamine release in the NAc core (Zhang et al., 2012; Exley et al., 2013). The same studies also found that oral nicotine self-administration decreased the influence of β2-containing nAChRs (Zhang et al., 2012) and α6-containing nAChRs (Exley et al., 2013) over dopamine release in the NAc core. Repeated nicotine also decreases α6β2* receptor control over dopamine release in the NAc shell and ventral putamen of nonhuman primates (Perez et al., 2009, 2012).

Our lab has recently shown individual variation in the degree to which nAChRs modulate dopamine release in the NAc core and that this variation correlates with a behavioral measure of vulnerability to high levels of early drug intake (Siciliano et al., 2017). Dopamine release in the NAc core is necessary for the incentive motivation of cues (Saunders & Robinson, 2012) and cue-induced drug seeking (Bossert et al., 2007; Saunders et al., 2013). Given the importance of nAChRs in modulating NAc dopamine release and their role in the dual-reinforcement effects of nicotine, chronic nicotine may modulate dopamine signals in a manner that drives further cue-induced drug seeking and use.

Additionally, it has been hypothesized that locomotor sensitization to nicotine is a marker of vulnerability to nicotine addiction (DiFranza & Wellman, 2007). Given our previous work, we were also interested in whether individual differences in nicotine-induced locomotor sensitization would correlate with nicotine-induced changes in nAChR modulation of dopamine release.

To examine the effects of chronic nicotine on nAChR modulation of dopamine release in the NAc core, we used *ex vivo* fast-scan cyclic voltammetry (FSCV) to measure dopamine release in rats following seven days of nicotine exposure. Various stimulation parameters were used to model a range of dopamine neuron firing patterns. Then, non-selective and selective nAChR antagonists were used to examine whether repeated nicotine altered nAChR modulation of NAc dopamine release. We then assessed whether magnitude of locomotor sensitization following repeated nicotine correlated with nicotine-induced modulation of dopamine release across tonic and phasic stimulations.

## Materials and Methods

### Animals

Adult male Sprague-Dawley rats (325-350 grams, Harlan Sprague Dawley, Inc., Madison, WI) were maintained on a 12:12 h reverse light/dark cycle (4:00 a.m. lights off; 4:00 p.m. lights on) with food and water available *ad libitum.* All animals were maintained according to the National Institutes of Health guidelines in Association for Assessment and Accreditation of Laboratory Animal Care accredited facilities. The experimental protocol was approved by the Institutional Animal Care and Use Committee at Wake Forest School of Medicine.

### Locomotor assessment and nicotine exposure

Rats were given at least a week to acclimate to the housing environment and light cycle prior to the start of experiments. All locomotor testing occurred during the dark/active cycle (9:00AM) to prevent sleep from contributing to variability in locomotor activity. Rats were first transferred to the dark locomotor testing room for one hour to habituate in their home cages. They were then placed in an acrylic locomotor activity chamber (45.7 cm x 45.7 cm x 30.4 cm) where their locomotor activity was monitored for 90 minutes. Rats were then subcutaneously injected on the flank with 0.9% saline solution, 0.2 mg/kg nicotine, or 0.4 mg/kg nicotine and replaced in the activity chamber for another 90 minutes. Nicotine (0.2-0.4 mg/kg, s.c.) or saline was administered for an additional six consecutive days during their active/dark cycle. On the seventh (last) day, locomotion was reassessed as on day one. Activity was recorded using Noldus^®^ video camera system and analyzed using EthoVision XT (version 11.5).

### Ex vivo fast scan cyclic voltammetry

The day after the final injection, rats were anesthetized with isoflurane and euthanized by decapitation. Brains were rapidly removed and transferred into ice-cold, pre-oxygenated (95% O_2_ / 5% CO_2_) artificial cerebral spinal fluid (aCSF) containing (in mM): NaCl (126), KCl (2.5), monobasic NaH_2_PO_4_ (1.2), CaCl_2_ (2.4), MgCl_2_ (1.2), NaHCO_3_ (25), dextrose (D-glucose) (11), and L-ascorbic acid (0.4). Tissue was sectioned into 400 μm-thick coronal striatal slices with a compresstome^®^ VF-300 vibrating microtome (Precisionary Instruments, San Jose, California). Brain slices were placed in submersion recording chambers and then perfused at 1mL/min at 32°C with oxygenated aCSF.

FSCV was used to assess dopamine release in the NAc core in rat brain slices (Figure 1A). A bipolar stimulating electrode was placed 100-150 μM from a carbon-fiber recording electrode (100-200 μm length, 7 µm diameter) in the NAc core (Figure 1A). Dopamine release was initially evoked by a single electrical pulse (750 μA, 2 msec, monophasic) applied to the tissue every 3 minutes.

**Figure 1.**
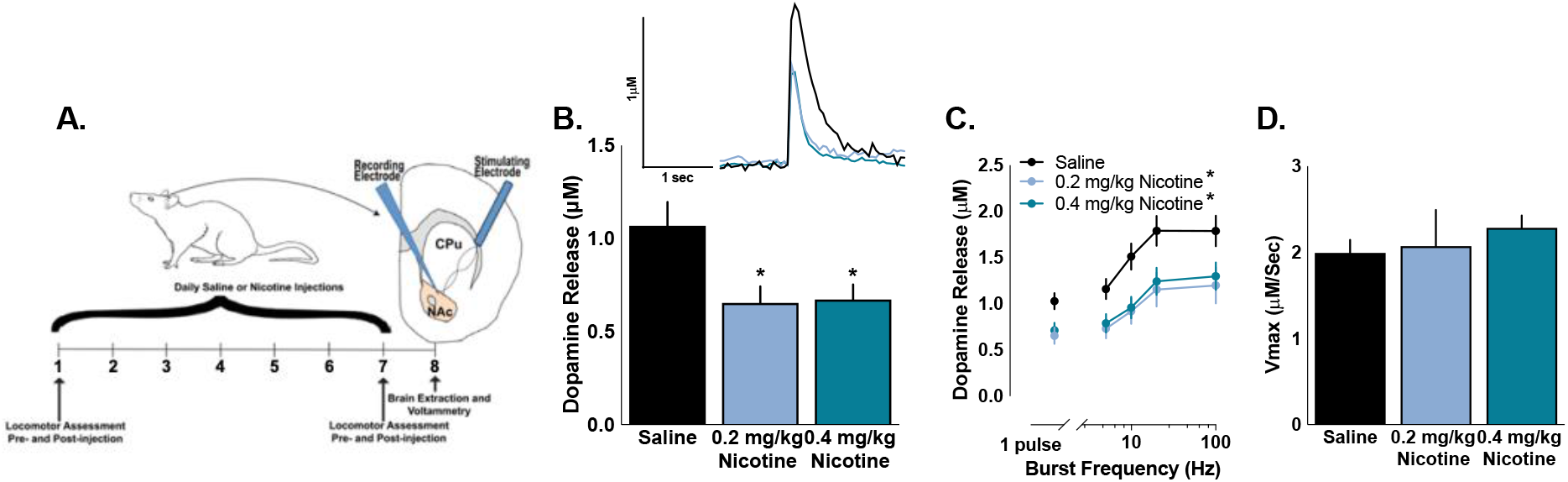
Chronic nicotine administration lowers dopamine signaling. (A) Experimental timeline of locomotor assessments, nicotine injections, and voltammetry. Rats were given subcutaneous injections of saline or 0.2 mg/kg or 0.4 mg/kg nicotine for seven days, with locomotor assessment on Days 1 and 7. On the eighth day, brains were extracted and *ex vivo* voltammetry was used to examine dopamine release in the nucleus accumbens core. (B) Chronic exposure to nicotine lowers electrically-stimulated single pulse dopamine release compared to saline. (C) Nicotine decreases both dopamine release, but does not differ between doses. (D) Maximal rate of diddopamine uptake (*V*_max_) is unaffected by nicotine exposure. Bars and symbols represent means ± SEMs, *p < 0.05.

Extracellular dopamine was recorded by applying a triangular waveform from −0.4 to 1.2 V and back to −0.4 (Ag vs AgCl) at a scan rate of 400 V/s using by a carbon fiber electrode. Voltammograms were recorded at the carbon fiber electrode every 100msec. Once dopamine response was stable (three consecutive collections with <10% variability), five-pulse stimulations were applied at varying burst frequencies (5, 10, 20, or 100 Hz) to model the physiological range of dopamine neuron firing. After assessing the dopamine response to single and multi-pulse stimulations, various compounds targeting nAChRs (nicotine, 500 nM; mecamylamine [MEC], 2 μM; dihydro-beta-erythroidine [DHβE], 500 nM; α-conotoxin MII [α-Ctx MII], 100 nM) were bath applied and dopamine response equilibrated (three collections within 10% variability) to single pulse stimulation. Separate slices from the same animal were used to test each drug independently, and the same frequency-response curves assessed under drug-free conditions were reassessed following drug application in each slice. In order to test the distinct contributions of α6* and non-α6* nAChRs, we added α-Ctx MII and DhβE in a cumulative fashion, equilibrating and testing single and multi-pulse frequencies (described above) following α-Ctx MII and then DhβE. Changes in dopamine signaling between α-Ctx MII [a selective α6 nAChR antagonist (Nicke et al., 2004)] alone and in combination with DHβE (a β2 nAChR antagonist) differentiated the contribution of α6* and non-α6* β2-containing nAChRs. We focused on nAChRs containing α6 subunits due to its role in modulating dopamine release in the NAc (Wickham et al., 2013; Siciliano et al., 2017).

Demon Voltammetry and Analysis software was used to acquire and model FSCV data (Yorgason et al., 2011). Recording electrodes were calibrated by recording electrical current responses (in nA) to a known concentration of dopamine (3 μM) using a flow-injection system. This was used to convert electrical current to dopamine concentration. Michaelis-Menten kinetics were used to determine maximal rate of dopamine uptake (*V*max) (Ferris et al., 2013).

### Statistical analysis

Single pulse dopamine release and *V*max were compared by one-way ANOVA and Tukey’s multiple comparison in case of significance. Dopamine release across multiple frequency stimulations was normalized to each subject’s pre-drug single pulse dopamine release. Multipulse dopamine release and locomotor activity were compared by two- or three-factor mixed design ANOVA. In the case of significant interactions, Bonferroni post-hoc comparisons were used. Percent changes in dopamine release following drug application were compared using two-tailed unpaired *t*-tests or two-factor mixed design ANOVA. Locomotor sensitization was assessed using one-sample *t*-tests against no change. Pearson product-moment correlation was used to assess the relationship between nicotine locomotor sensitization and nicotine- and MEC-induced modulation of dopamine release. All statistics were performed using GraphPad Prism 7 (Graphpad Software, La Jolla, CA) or SPSS v. 24 (International Business Machine Corporation, Armonk, NY) with α≤ 0.05. Values >2 standard deviations above or below the mean were considered outliers and excluded. Data are presented as mean ± SEM.

## Results

### Repeated nicotine exposure decreases dopamine release in the NAc core

We first examined whether repeated exposure to nicotine altered dopamine release in the NAc core. Rats were exposed to nicotine (0.2 or 0.4 mg/kg, s.c.) for seven consecutive days, then *ex vivo* voltammetry was used to assess dopamine release in the NAc core the day after the final injection of nicotine (Fig. 1A). Repeated injections of nicotine significantly decreased the magnitude of dopamine evoked by a single pulse (*F*_2, 45_ = 5.058, p = 0.01), with no significant difference between the doses of nicotine (Fig. 1B). Dopamine release was elicited by five pulse stimulations across the range of physiological dopamine neuron firing in order to examine dopamine signaling at frequencies that model tonic- and phasic-like firing patterns. As expected, frequency of the five pulse stimulation modulates dopamine release (*F*_4, 184_ = 92.86, p < 0.001), with higher frequencies increasing dopamine release. In concurrence with the single pulse dopamine release, repeated nicotine decreased dopamine release across the range of frequencies (*F*_2, 46_ = 4.964, p = 0.011) and the decrease in dopamine release was not different between the doses of nicotine (Fig. 1C). The maximal rate of dopamine uptake (*V*max) was not impacted by repeated exposure to nicotine (*F*_2, 41_ = 0.528, p = 0.594)(Fig. 1D). Given that the dose of nicotine did not differentially impact the magnitude of decrease in dopamine release, we focused only on the 0.4 mg/kg dose of nicotine in subsequent experiments.

### α6-containing nAChR regulation of dopamine release is altered following repeated nicotine

To examine whether repeated exposure to nicotine had functional consequences on nAChR modulation of dopamine release in the NAc core, we assessed dopamine release across a range of frequencies following bath application of drugs that target nAChRs. Reductions in dopamine release to single pulse stimulations, in particular, could be attributed to reductions in acetylcholine (ACh) facilitation of dopamine release magnitude. Striatal cholinergic interneurons (CIN) increase dopamine release in the NAc core by activating nAChRs on dopamine terminals (Cachope et al., 2012; Threlfell et al., 2012; Brimblecombe et al., 2018) and antagonism or desensitization of nAChRs decreases single pulse dopamine release (Rice & Cragg, 2004). Surprisingly, a history of nicotine exposure did not alter the magnitude of decrease in dopamine release to single pulse stimulations following a desensitizing dose of nicotine (*t*_32_ = 0.098, p = 0.922) (Fig. 2A) or MEC (a non-selective nAChR antagonist) (*t*_12_ = 0.87, p = 0.401) (Fig. 2B).

**Figure 2.**
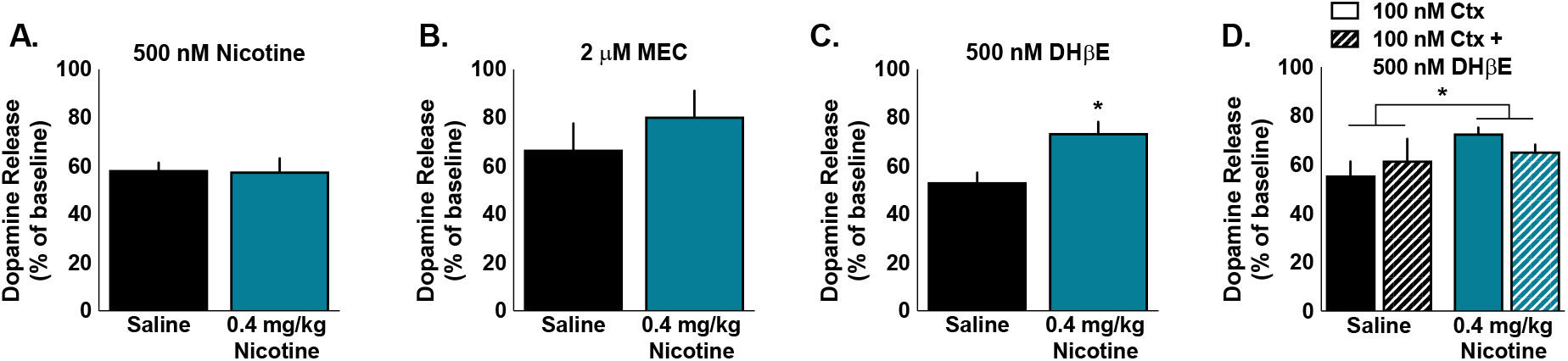
Chronic nicotine alters α6β2 nAChR modulation of single pulse dopamine. (A) Chronic nicotine exposure did not alter the effect of nicotine (500 nM) or (B) MEC [a non-selective nAChR antagonist (2 μM)] on single pulse dopamine release in the NAc core. (C) DHβE [a selective β2 nAChR antagonist (500 nM)] decreased dopamine release significantly more in saline than nicotine treated rats. (D) Chronic nicotine exposure also blunted the decrease in single pulse dopamine release following application of α-Ctx MII [a selective α6 nAChR antagonist (100 nM)] followed by DHβE. This order was used to differentiate the effect of α6 and non-α6 nAChRs. Bars and symbols represent means ± SEMs, *p < 0.05.

We next used selective nAChR antagonists to examine whether β2* nAChR-modulation of dopamine was altered by repeated exposure to nicotine since β2-containing nAChRs are necessary for the reinforcing effects of nicotine and for nicotine-induced increases in NAc dopamine (Picciotto et al., 1998; Maskos et al., 2005). To determine this, we examined dopamine release following a bath application of DHβE [a β2-selective antagonist (500 nM)]. A history of nicotine exposure significantly blunted the decreasing effects of DHβE on single pulse dopamine release (*t*_26_ = 3.269, p = 0.003)(Fig. 2C). To examine the contribution of α6* and non-α6* β2-containing nAChRs to the changes in β2-containing nAChR modulation of dopamine, we applied α-Ctx MII [a selective α6 antagonist (100 nM)] followed by DHβE. Consistent with the DHβE results above, saline treated animals had a significantly greater decrease of single dose dopamine release following treatment with α-Ctx MII alone (solid bars) and α-Ctx MII + DHβE (main effect of group: *F*_1, 10_ = 8.358, p = 0.016). DHβE did not significantly modulate the effect of α-Ctx MII on dopamine release in either group (main effect of drug: *F*_1, 10_ = 0.012, p = 0.914; interaction group*drug: *F*_1, 10_ = 1.293, p = 0.282).

Since nAChRs modulate dopamine release in a frequency dependent manner and the effects of cholinergic and nicotine-induced modulation of dopamine on behavior are hypothesized to be mediated by frequency-dependent gating of dopamine (Rice & Cragg, 2004), we wanted to examine the effects of nicotine and non-selective and selective nAChR antagonists across a range of physiologically relevant frequencies in the NAc core. To determine this, we used five pulse stimulations across a range of frequencies to model tonic- and phasic-like firing patterns before and after nicotine or nAChR antagonism. As expected, nicotine modulated dopamine release in a frequency-dependent manner, decreasing dopamine release to single pulse and low frequency stimulation, but not to the highest frequency stimulation (interaction drug*frequency: *F*_4, 124_ = 15.383, p < 0.001). However, repeated nicotine exposure did not change nicotine-induced modulation of dopamine release (main effect of group: *F*_1, 31_ = 0.026, p = 0.874)(Fig. 3A). Similar to the effects of nicotine, MEC decreased single pulse and low frequency dopamine release, but did not affect high frequency dopamine release, and this modulation was not changed by a history of nicotine exposure (main effect of group: *F*_1, 13_ = 0.001, p = 0.982; interaction drug*frequency: *F*_4, 52_ = 5.419, p = 0.001)(Fig. 3B).

**Figure 3.**
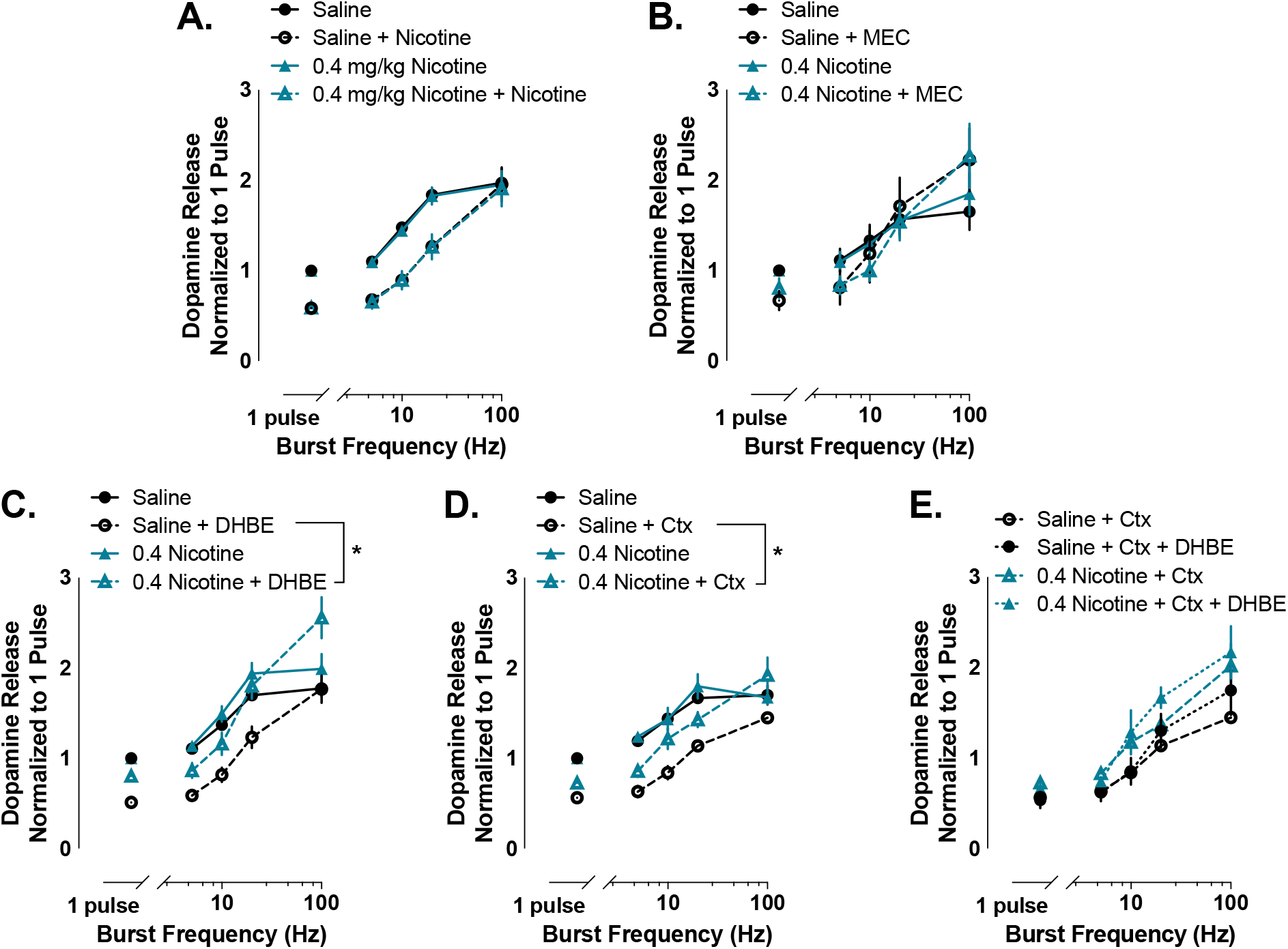
α6 nAChR modulation of dopamine release is altered following chronic nicotine. (A) Nicotine (500 nM) decreased dopamine release to single pulse and low frequency stimulation, but not the highest stimulation frequency. Chronic nicotine exposure did not change nicotine-induced modulation of dopamine release. (B) MEC (2 μM) decreased dopamine release to single pulse and low frequency stimulation in both saline and nicotine treated animals. (C) DHβE (500 nM) and (D) α-Ctx MII (100 nM) also modulate dopamine release in a frequency-dependent manner, but dopamine release is higher in rats with chronic nicotine exposure and shows facilitation at higher frequencies. (E) The application of DHβE following α-Ctx MII does not significantly change dopamine release. Bars and symbols represent means ± SEMs, *p < 0.05. Note: Not all significant interactions are visually represented.

To determine the role of β2-containing nAChRs and isolate the role of α6 and non-α6 containing β2* nAChRs on dopamine release to tonic and phasic firing patterns, we applied DHβE alone and following application of α-Ctx MII. As with nicotine, DHβE decreased dopamine to single pulse and low frequency stimulations, but not to the highest frequency stimulations (interaction drug*frequency: *F*_4, 104_ = 22.468, p < 0.001). Interestingly, DHβE decreased dopamine release across the range of stimulation frequencies significantly more in saline treated rats than rats with repeated nicotine exposure (interaction drug*group: *F*_1, 26_ = 7.608, p = 0.011) (Fig. 3C). In agreement with this effect being driven by α6-containing nAChRs, dopamine release was significantly higher in rats with repeated nicotine exposure following α-Ctx MII application (interaction drug*group: *F*_1, 10_ = 4.905, p = 0.05) (Fig. 3D). Additionally, DHβE did not significantly affect dopamine release in saline or nicotine treated rats when applied after α-Ctx MII (main effect of drug: *F*_1, 8_ = 0.739, p = 0.415), although rats with repeated nicotine exposure did have higher dopamine release than saline treated rats at the two highest frequency stimulations following α-Ctx MII and subsequent DHβE (interaction group*frequency: *F*_4, 32_ = 4.148, p = 0.008) (Fig. 3E).

### The magnitude of nicotine sensitization predicts nicotine-induced modulation of striatal dopamine release at phasic firing frequencies

Repeated exposure to nicotine increases nicotine-induced locomotion and nicotine sensitization is hypothesized to be a marker of vulnerability to nicotine addiction (DiFranza & Wellman, 2007). Previous work from our lab has shown that nAChR modulation of dopamine release in the NAc core correlates with another model of vulnerability (high and low responders) (Siciliano et al., 2017). Because of these findings, we were interested in whether nAChR modulation of dopamine release may predict locomotor sensitization following repeated nicotine exposure. Acute nicotine and repeated saline did not affect locomotion. As expected, repeated nicotine injections increased locomotion following a nicotine injection, but did not alter baseline locomotion (interaction drug*time*day: *F*_20, 1020_ = 4.994, p < 0.001) (Fig. 4A). Repeated injections of nicotine significantly increased nicotine-induced locomotion (one-sample t-test: *t*_31_ = 7.31, p < 0.001), while repeated saline did not alter locomotion following a saline injection (one sample t-test: *t*_21_ = 0.881, p = 0.388) (Fig. 4A).

**Figure 4.**
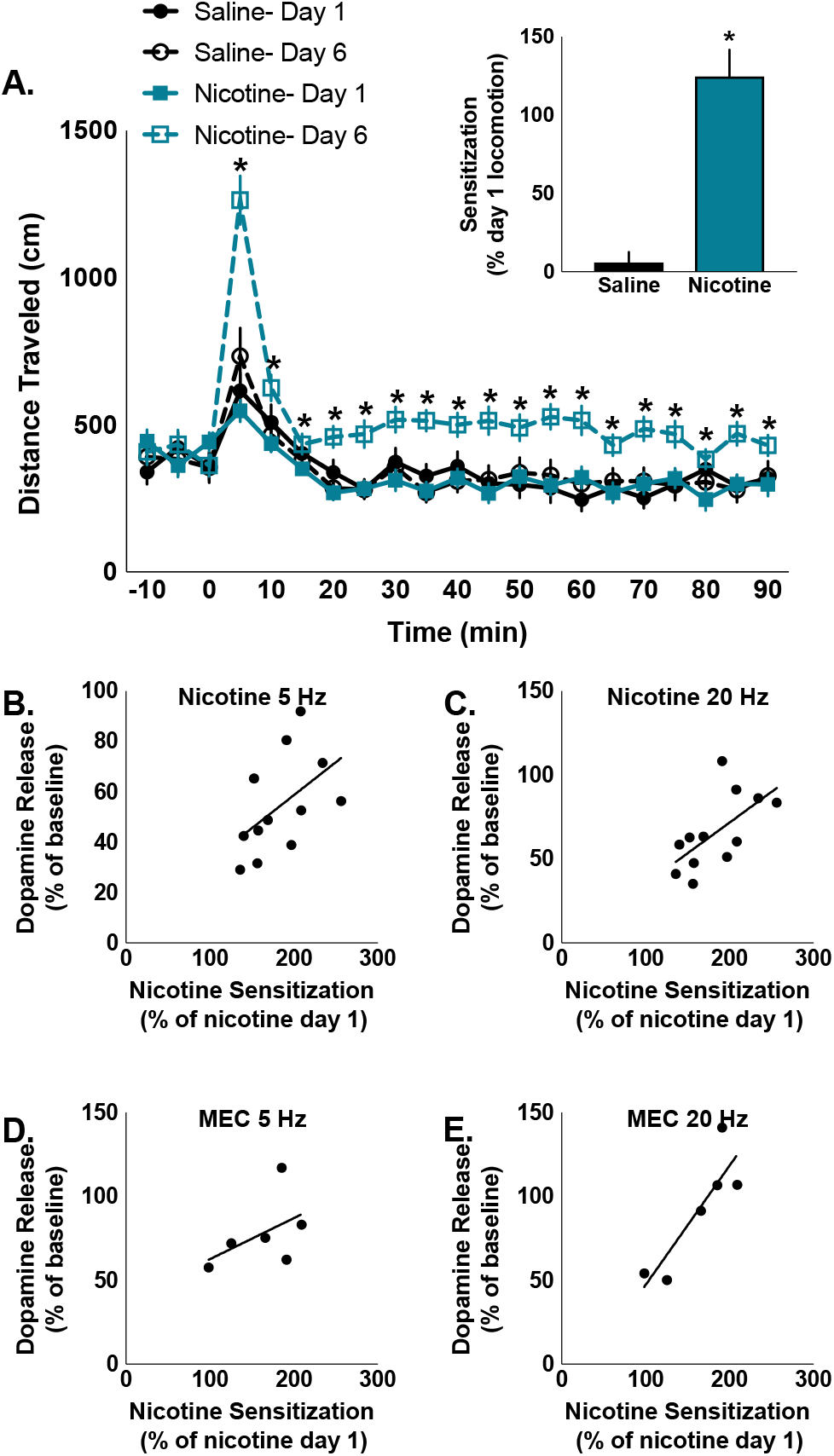
The magnitude of nicotine-induced sensitization predicts the effects of nicotine on dopamine release to phasic firing in the NAc. (A) Nicotine increases post-injection locomotion significantly more following repeated nicotine exposure. Repeated saline and acute nicotine do not alter locomotion. Inset: Repeated injections of nicotine significantly changed nicotine-induced locomotion, while repeated injections of saline did not change locomotion following a saline injection. (B) Magnitude of nicotine-induced locomotor sensitization is not predicted by changes in tonic (5 Hz) stimulations following bath application of nicotine (500 nM), (C) but is predicted by nicotine-induced changes to dopamine release following phasic (20 Hz) stimulation. (D) Similarly, magnitude of nicotine locomotor sensitization was not correlated with MEC-induced (2 μM) changes in dopamine release to tonic stimulation, (E) but did positively correlate with changes to phasic dopamine release following MEC application. Bars and symbols represent means ± SEMs, *p < 0.05.

There was individual variation in how much repeated nicotine sensitized nicotine-induced locomotion. We examined whether locomotor sensitization to nicotine predicted the effects of nicotine on dopamine release in the NAc core. Locomotor sensitization to nicotine did not correlate with the effects of nicotine on tonic frequency stimulation of dopamine (5 Hz: r = 0.514, p = 0.088) (Fig. 4B), but did correlate with nicotine-induced modulation of dopamine following phasic firing rates (20 Hz: r = 0.632, p = 0.028) (Fig. 4C). In agreement with the effects of nicotine, locomotor sensitization did not correlate with MEC-induced modulation of dopamine release to tonic firing rates (5 Hz: r = 0.488, p = 0.326) (Fig. 4D), but did correlate with dopamine release to phasic firing stimulation (20 Hz: r = 0.87, p = 0.024) (Fig. 4E).

## Discussion

Our study demonstrates that seven consecutive days of nicotine injections in rats significantly reduced dopamine release magnitude in the NAc core following single pulse electrical stimulation and five pulse stimulations at frequencies ranging from 5 Hz to 100 Hz. This reduction was observed after repeated exposure to both 0.2 and 0.4 mg/kg nicotine. The change in dopamine release was not associated with a change in dopamine transporter function, indicated by no change in the maximal rate of dopamine uptake (Vmax) across groups. Further examination of functional changes in nAChR modulation of dopamine release revealed that repeated nicotine blunted reductions in dopamine release elicited by the β2 selective antagonist, DHβE, and the α6 selective antagonist, α-Ctx MII. This blunted effect of selective antagonists was present across the entire range of stimulation frequencies examined. For higher frequencies, the repeated nicotine injections altered the dopamine response to nAChR antagonists from no change (saline) to a facilitation of dopamine release following nAChR blockade. Finally, repeated nicotine sensitized locomotor response to a systemic nicotine challenge. The magnitude of sensitization was predicted by the degree to which blocking nAChRs modulated dopamine release magnitude to frequencies that model phasic (20 Hz), but not tonic (5 Hz), firing of dopamine neurons.

Our results indicate that repeated nicotine administration (0.2 and 0.4 mg/kg) decreases dopamine release during both single pulse stimulation and multiple pulse stimulations across a range of frequencies that model tonic and phasic firing of dopamine neurons. Moreover, repeated nicotine did not alter the frequency dependent nature of dopamine release magnitude. Although no previous study has assessed dopamine release following nicotine exposure using a model of repeated nicotine injections [which more closely mimics the rapid rate of nicotine delivery and pharmacokinetics seen with smoking than oral administration (Benowitz et al., 2009; Perez et al., 2012; Exley et al., 2013; Koranda et al., 2014)] or studied rats, these results are consistent with previous published reports showing decreases in dopamine release in the NAc and caudate-putamen in mice following several weeks of oral nicotine treatment (Zhang et al., 2012; Exley et al., 2013) and the NAc shell and ventral putamen of squirrel monkeys following several months of oral nicotine treatment (Perez et al., 2009, 2012). Our model of seven days of repeated nicotine injections in rats produced a magnitude of dopamine reduction more similar to the effect sizes in monkeys compared to the studies in mice. Additionally, our effects were not due to differences in dopamine uptake through the dopamine transporter since Vmax was not changed by nicotine exposure.

We next examined the degree to which these nicotine-induced reductions in dopamine release could be attributed to reductions in acetylcholine (ACh) facilitation of dopamine release magnitude. Previous work has established that striatal cholinergic interneurons (CIN) supply abundant ACh to the NAc, which can facilitate dopamine release magnitude via nAChRs located directly on dopamine axons. Indeed, acute β2* blockade or desensitization lowers the probability of dopamine release in response to single pulse and multiple pulse stimulations that model tonic firing of dopamine neurons (Rice & Cragg, 2004; Zhang & Sulzer, 2004; Zhang et al., 2009). α6β2-containing nAChRs dominate ACh influence over dopamine release in the ventral striatum / NAc core, while α5-containing nAChRs play a larger role in the dorsal striatum (Exley et al., 2008). Thus, we hypothesized that if ACh release from CINs was blunted (or has less influence over dopamine release) in the NAc core following repeated nicotine injections, then both β2 and α6 selective antagonists would be less effective at reducing dopamine release in nicotine treated rats compared to saline treated rats. Consistent with this hypothesis, both the β2 selective antagonist, DHβE, and the α6 selective antagonist, α-Ctx MII, were less effective at reducing dopamine release in animals treated with nicotine. The nonselective antagonist, mecamylamine, and a desensitizing dose of nicotine decreased dopamine equally in both saline and repeated nicotine groups. The difference in outcome between these nonselective (or desensitizing) compounds and the α6β2 selective compounds is unclear. One major difference between these classes of drugs is that the nonselective compounds also bind α7 nAChRs, which are located on glutamate afferents in our slice preparation and could effectively reduce excitatory drive onto dopamine axons when blocked or desensitized with mecamylamine or nicotine, respectively. A reduction in excitatory drive on dopamine axons has the potential to decrease dopamine release to floor effects and mask the α6β2* mediated effects observed using our selective antagonists. Regardless, the involvement of α6β2* nAChRs is consistent with previous voltammetric studies that showed α6β2-containing receptors are primarily responsible for nAChR-evoked dopamine release in the ventral striatum. This is further supported by the fact that DHβE had no effect on dopamine release magnitude when administered after α-Ctx MII, suggesting minimal contribution from non-α6 containing nAChRs (Exley et al., 2008; Siciliano et al., 2017) to our nicotine treatment differences in dopamine release.

We next examined whether repeated nicotine injections sensitized locomotor response to nicotine challenge, as previously reported (see DiFranza & Wellman, 2007), and whether the degree to which sensitized locomotor activity relates to the magnitude of nicotine’s effect on NAc phasic signals in a slice. We show that a seven day regimen of once daily nicotine injections (0.4 mg/kg, s.c.) sensitizes locomotor activity, with nicotine treated rats more than doubling their locomotor activity after the sixth nicotine injection compared to the first. Although elevations in locomotor activity are the most robust ≤ 15 minutes post injection, elevations are sustained through the entire session. We also found that the magnitude of nicotine-induced locomotor sensitization is not predicted by changes in tonic (5 Hz) stimulations following bath application of either nicotine or MEC, but did positively correlate with changes to phasic (20 Hz) dopamine release with both nicotine and MEC. This data is particularly interesting given our previous data that phasic, but not tonic, dopamine release magnitude following bath application of nicotine and MEC correlates with locomotor response to novelty (Siciliano et al., 2017), a strong predictor of acquisition rates for several drugs of abuse (Piazza et al., 1989; Mantsch et al., 2001; Ferris et al., 2013). Thus, nicotine modulation of NAc core phasic dopamine release correlates with two markers of vulnerability to substance use disorders: one in drug naïve animals and one following repeated drug exposure. This generality suggests that striatal nAChR modulation of NAc core dopamine (or the interaction of striatal acetylcholine and dopamine) may be a potential biomarker of vulnerability to SUD, or directly mediate SUD vulnerability. Indeed, recent work has shown mechanistic links between ACh signaling through nAChRs on dopamine axons and modulation of cue-induced motivation for natural rewards (Collins et al., 2019). Future studies will need to explore whether such findings extend to drug seeking.

In conclusion, we found that repeated nicotine injections blunt dopamine release equally across a range of stimulation frequencies that model both tonic and phasic firing of dopamine neurons and that repeated nicotine decreased the ability of α6β2-containing nAChRs to modulate dopamine release. This deficit in dopamine function may underlie, in part, increased vulnerability to nicotine use following repeated exposure to nicotine. In particular, CIN modulation of dopamine release (mediated through nAChRs) is thought to be essential for reward-related learning (Cragg, 2006) and dysregulation of this system may alter responses to rewards (i.e., nicotine) and reward-related cues in a manner that drive nicotine use disorder. We also extended our earlier work on the relationships between locomotor response to either novelty or acute nicotine and dopamine release magnitude following nicotine administration in the NAc. Indeed, we found that nicotine locomotor sensitization, a potential marker of vulnerability to nicotine dependence, correlates with nicotine and MEC modulation of phasic dopamine release. Together, these data suggest that repeated nicotine exposure alters nAChR control over dopamine release in the NAc core in a manner that is consistent with changes that may serve as a biomarker for vulnerability to nicotine use, or a mechanism for such vulnerability.

